# Systematic screen of PKR reveals genetic variants that broadly evade divergent viral pseudosubstrate inhibitors

**DOI:** 10.64898/2026.08.11.744216

**Authors:** Michael J. Chambers, Tristan R. Grieve, Sophia B. Scobell, Meru J. Sadhu

## Abstract

Evolutionary arms races can arise at the contact surfaces between host and viral proteins, producing dynamic spaces in which genetic variants are continually pursued even while protein function must be maintained. A striking case is protein kinase R (PKR), an innate immune protein that detects viral replication and halts protein synthesis by phosphorylating the initiation factor eIF2α. PKR is targeted by many viral antagonists, including pseudosubstrate inhibitors encoded by poxviruses and ranaviruses that mimic eIF2α. We previously found that the eIF2α-binding surface of human PKR is highly malleable against the vaccinia virus pseudosubstrate inhibitor K3. Here, we screen our library of SNP-accessible PKR variants against five viral inhibitors of increasing divergence: K3 orthologs from vaccinia, variola, tanapox, and myxoma viruses, and the independently derived vIF2α from *Rana catesbeiana* virus Z. Resistance-conferring variants are readily accessible against every inhibitor tested and are often shared across phylogenetically diverse K3 orthologs and the ranavirus inhibitor, suggesting that PKR escape variants can exploit features common to pseudosubstrate inhibitors. Many variants beneficial against multiple inhibitors correspond to sites under positive selection across vertebrates. Inhibitor-specific effects are largely explained by differences in contact residues between PKR and each inhibitor. Notably, no PKR variant became newly susceptible to myxoma K3, which does not naturally inhibit human PKR. The eIF2α-binding surface of PKR is thus broadly navigable against genetically diverse viral pseudosubstrate inhibitors, potentiating its evolutionary capacity to combat viral inhibition without incurring new vulnerabilities.

## INTRODUCTION

The innate immune system provides the host with a first line of defense against infectious agents. Protein kinase R (PKR), a component of the innate immune system, is activated by binding viral intracellular double-stranded RNA. Active PKR phosphorylates the translation initiation factor eIF2α to halt translation, preventing viral protein synthesis and halting viral replication. Viruses employ a variety of strategies to block or circumvent PKR antiviral activity. One strategy is to encode mimics of eIF2α, replicating the OB-fold domain that interfaces with the PKR kinase domain but blocks phosphorylation of eIF2α. Many poxviruses encode eIF2α mimics, and the breadth of poxvirus species in which they are present suggests they first emerged at least 160,000 years ago^2^, in which time they have diverged substantially (Fig. 1A). Humans have a long history of interaction with poxviruses, including variola virus, the causative agent of smallpox disease; monkeypox virus, which causes mpox disease; and vaccinia virus, which is used in live-virus vaccination against both smallpox and mpox. Many other poxvirus species can infect humans, including hundreds of known cases of infection by tanapox virus^3,4^. The vaccinia K3 protein is the best-studied viral eIF2α mimic. The variola K3 ortholog C3, hereafter referred to as variola K3, is approximately 80% identical to vaccinia K3, while the tanapox K3 ortholog is approximately 40% identical (Fig. 1A, Extended Data Fig. 1). Monkeypox virus is a close relative of vaccinia virus ^5^ but it lacks a K3 ortholog due to the gain of a premature stop codon. Outside of poxviruses, eIF2α mimicry emerged independently in ranaviruses, which infect fishes, amphibians, and reptiles^6^

**Fig. 1.**
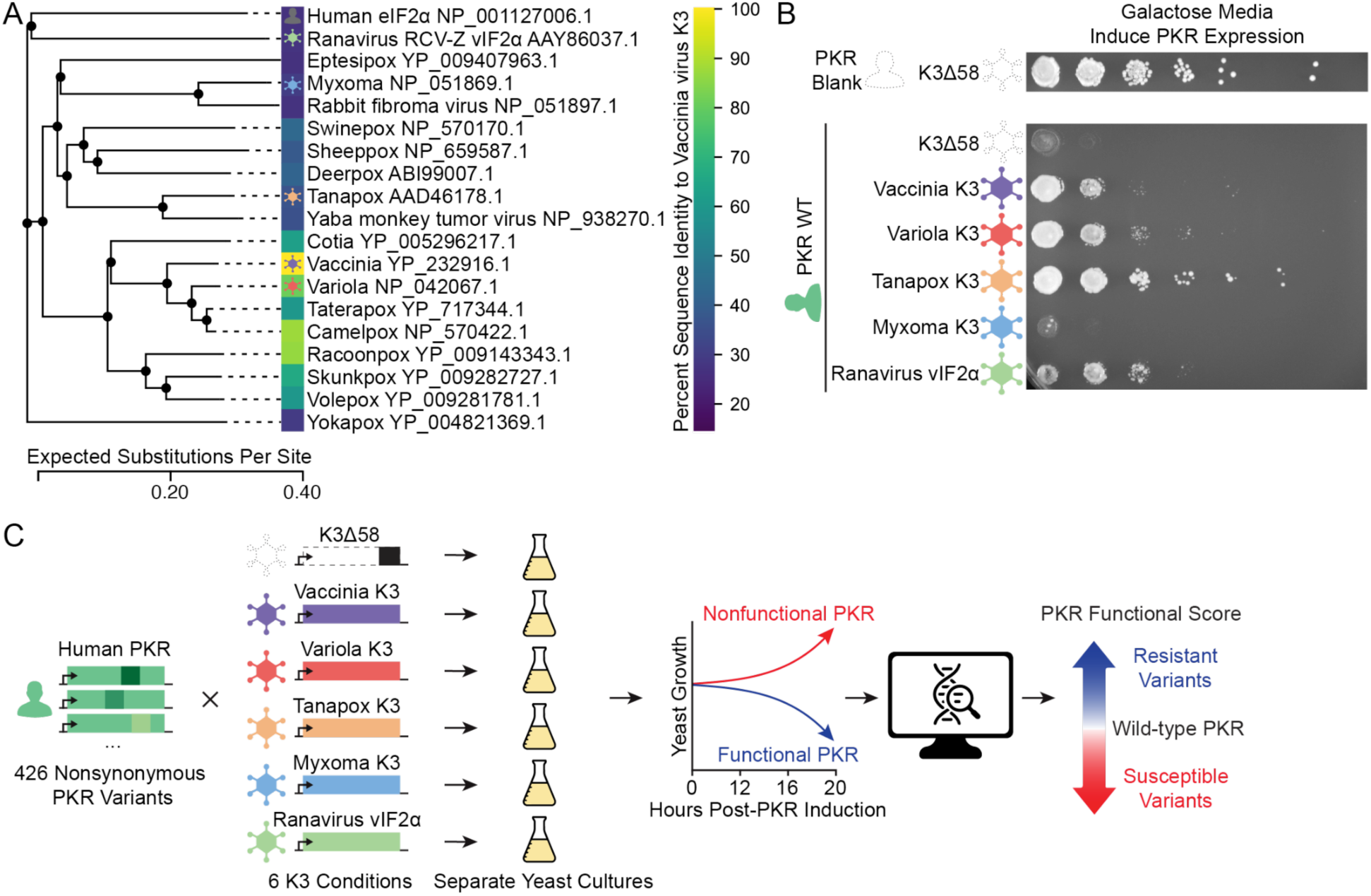
Viral mimics of eIF2α. (A) Phylogenetic tree of eIF2α homologs, including human eIF2α, the viral vIF2α from Ranavirus RCV-Z, and poxviral eIF2α homologs. Branch length corresponds to substitutions per site. Tree is midpoint rooted for visualization purposes. A heatmap of amino acid percent identity to vaccinia virus is shown to the right of the tree. Five viral homologs chosen for characterization in this study are marked with virus symbols. The coloring of the virus symbols will be maintained throughout the figures. (B) PKR expression is toxic to the budding yeast *Saccharomyces cerevisiae*, and can be suppressed by viral eIF2α homologs. Yeast were serially diluted on galactose plates, inducing expression of human PKR from a *GAL1/CYC* promoter. Co-expression of viral eIF2α homologs from vaccinia, variola, tanapox, and Ranavirus allowed substantially improved yeast growth relative to expression of a nonfunctional mutant vaccinia K3 or the myxoma K3. (C) Methodological approach to explore the effects of variants of human PKR in the presence of different K3 orthologs. 426 nonsynonymous variants of PKR were generated and screened against K3Δ58, vaccinia K3, variola K3, tanapox K3, myxoma K3, and ranavirus vIF2α. Variant effects were characterized using a high-throughput yeast growth assay and massively parallel sequencing.

PKR has significant sequence diversity across species, with many sites having gained nonsynonymous changes at a higher than neutral rate ^7–9^, consistent with positive selection to escape viral inhibitors^10,11^. Several of these sites localize at or near the surface of PKR that is bound by eIF2α and poxvirus K3 orthologs. To examine the ability of PKR variants to disrupt K3 inhibition while maintaining eIF2α binding, we previously comprehensively tested the consequences of SNP-accessible variants in human PKR at its interaction surface shared by eIF2α and K3^12^ We found that PKR’s kinase activity towards eIF2α was highly tolerant of mutation, and yet many mutations disrupted vaccinia K3 inhibition. Overall, we found that human PKR has a favorable evolutionary landscape when challenged by vaccinia K3.

Here, we expanded on those results by characterizing PKR’s evolutionary landscape against four additional viral pseudosubstrate inhibitors that target PKR, encoded by the poxviruses variola virus^7^, tanapox virus^13^, and myxoma virus^14^; as well as ranavirus *Rana catesbeiana* virus Z (RCV-Z)^6^. The poxvirus K3 orthologs span 160,000 years of sequence diversity^15^, which allowed us to test PKR variants against inhibitors of varying levels of divergence. The myxoma K3 ortholog is known to be a poor inhibitor of human PKR^14^ and provided an opportunity to identify PKR variants that become newly susceptible to myxoma K3 inhibition. In contrast, RCV-Z independently derived an eIF2α mimic. This allowed us to differentiate whether PKR evades a unique feature of poxvirus pseudosubstrate inhibitors or if its evasion is a more general phenomenon across viral inhibitors. For all eIF2α mimics tested, we found that PKR could take many evolutionary routes to evasion. Moreover, the same variants often proved effective to evade multiple eIF2α mimics, which would be a useful feature in the evolutionary landscape of an innate immune protein faced with multiple viral antagonists. We also found variants that provided benefit against some viral inhibitors and not others, which allowed us to dissect the molecular basis of host-pathogen protein-protein compatibility. Finally, we found that no mutations in human PKR caused it to become susceptible to the rabbit-specific myxoma virus K3 ^14^. Overall, our expanded screen indicates that there are many evolutionary routes available for PKR to evade genetically diverse viral pseudosubstrate inhibitors.

## RESULTS

We previously made a library of mutations in the kinase domain of human PKR ^12^ focusing on mutations at sites predicted to be within 5 Å of eIF2α or K3 and sites under positive selection across vertebrate PKR homologs^7^ The 426 variants in our library are within a single nucleotide polymorphism relative to the reference human PKR DNA sequence (GenBank M85294.1:31–1686), as these are the variants most likely to be accessed in PKR’s evolutionary landscape. Variant PKR alleles can be characterized by expressing them in *Saccharomyces cerevisiae*, as functional PKR autoactivates and arrests growth via phosphorylation of yeast eIF2α^16,17^ Expression of viral eIF2α mimics that successfully inhibit PKR’s phosphorylation of eIF2α in this system suppress the growth arrest, allowing the identification of PKR mutations that evade viral inhibitors via their continued toxicity^7,8,18,19^. We previously developed a high-throughput version of this approach to characterize our library of PKR variants in the presence or absence of vaccinia K3^12^. Here, we extended our study to other viral inhibitors, namely the K3 orthologs present in variola virus, tanapox virus, and myxoma virus, as well as the independently derived eIF2α mimic from RCV-Z, hereafter referred to as ranavirus vIF2α. Aside from myxoma K3, which is known to be specific to rabbit PKR ^14^, these viral proteins all reduced the toxicity of human PKR in yeast (Fig. 1B) with strengths consistent with past results^7,13,18^. We characterized the ability of our PKR mutation library to evade these inhibitors with the high-throughput yeast assay (Fig. 1C). We additionally repeated our screen of PKR variants in the absence of any inhibitor (Extended Data Fig. 2) and in the presence of Vaccinia K3 (Extended Data Fig. 3), and observed strong correlations (Pearson correlation coefficient > 0.97) between our new results and our previously published findings (Extended Data Fig. 4). Any comparisons hereafter to the effect of PKR mutations in the presence or absence of vaccinia K3 will be to the most recent dataset. As in our previous study, we calculated a functional score for each PKR variant against each inhibitor: PKR variants that suppress yeast growth by evading the pseudosubstrate have high functional scores, and those that allow yeast growth, such as by being inhibited by the pseudosubstrate or otherwise unable to phosphorylate eIF2α, have low functional scores. We observed that nonsense variants of PKR had lower functionality than wild-type PKR in the presence of all the viral inhibitors (Extended Data Fig. 5), indicating that none of the inhibitors completely abolished PKR activity. Thus, yeast growth can be used to detect PKR variants that either improved or decreased evasion of variola K3, tanapox K3, and ranavirus vIF2α, as well as variants that confer susceptibility to myxoma K3.

### Variola K3

Variola virus, like vaccinia virus, belongs to the Orthopoxvirus genus in the Poxviridae family^20^. Variola virus is the causative agent of smallpox and has a strictly limited host range to humans; its last common ancestor with vaccinia virus existed approximately 4,000 years ago^5^. Variola K3 is 88 residues in length and is 80% identical to the sequence of vaccinia K3 (71 identical residues, Fig. 2A). 14 of these differing residues are within 5 Å of PKR and can be considered interfacing residues.

**Fig. 2.**
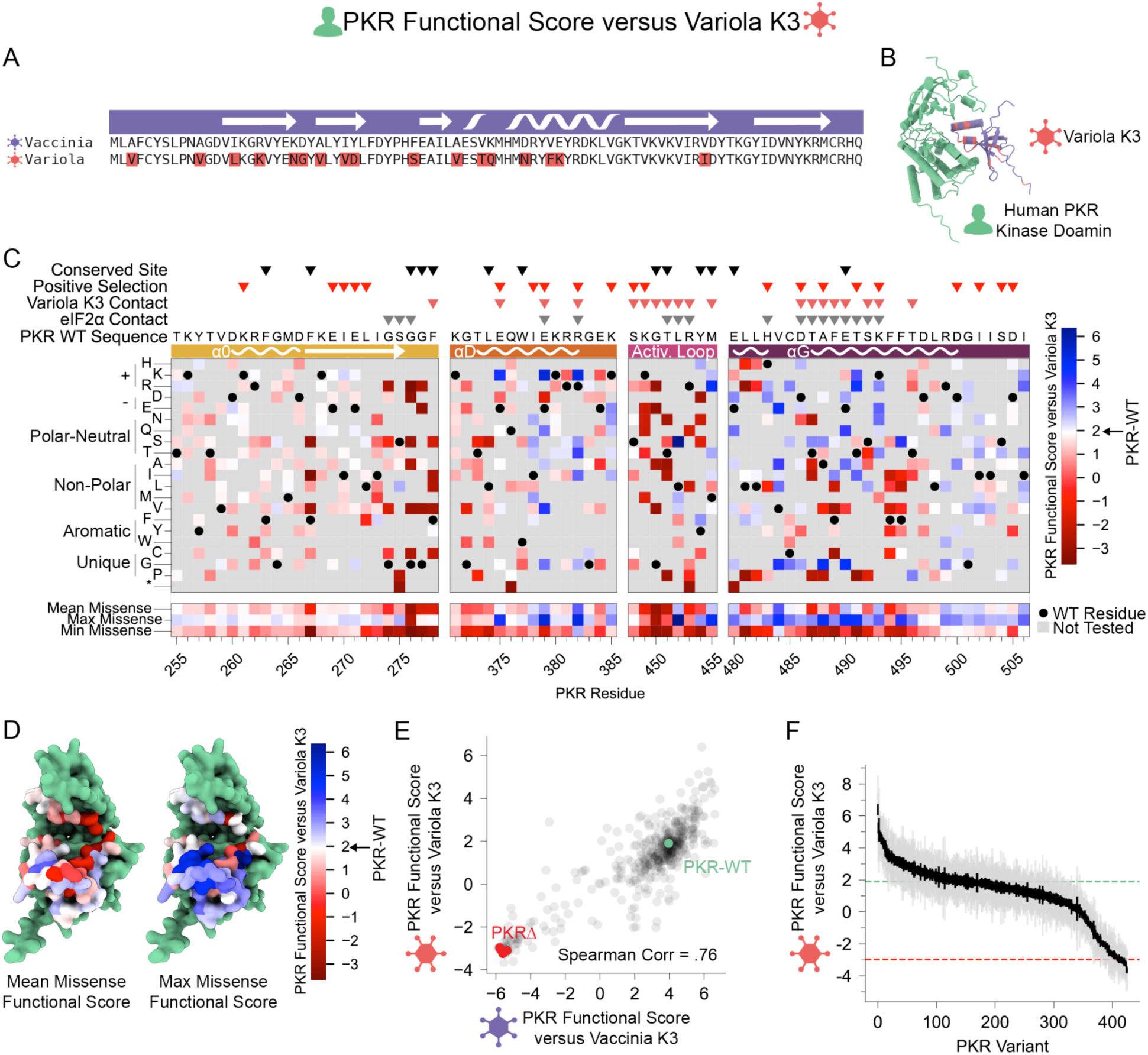
Many beneficial PKR variants are available against variola K3. (A) Alignment of the vaccinia and variola K3 homologs. The top track in purple shows secondary structure elements of vaccinia K3 (PDB 1LUZ)^33^. Amino acids differing in variola from vaccinia are highlighted in red. (B) Prediction of variola K3 bound to human PKR (green), generated by AlphaFold2-Multimer. Variant sites between variola and vaccinia K3 are colored red, while conserved sites are colored purple. (C) PKR functional scores versus variola K3 are colored ranging from susceptible (red) to WT-like (white) to resistant (blue). Wild-type PKR residues are denoted with black circles. Variants not accessible through a single base change are colored in gray, as they were not characterized. Sites conserved in ≥90% of vertebrate PKR homologs or that experienced positive selection^7^ are labelled with black and red triangles, respectively^7^ Contact sites are labelled based on AlphaFold2-Multimer prediction of being within 5 angstroms of the binding partner. (D) Surface residues of the PKR kinase domain colored by their mean (left) or maximum (right) PKR functional scores against variola K3, using identical coloring to panel C. (E) Correlation between PKR variant functional scores versus vaccinia and variola K3. The data point for wild-type PKR is colored green and the four data points for nonsense variants of PKR are colored red. Spearman correlation of 0.76 was calculated across all PKR variants. (F) PKR functional scores versus variola K3, ordered by function from high to low. Gray and black lines indicate the standard deviation and standard error, respectively, in the score across the barcoded replicates of each variant. The functional score of wild-type PKR is denoted with a horizontal green dashed line, and the average functional score of the four nonsense PKR variants is denoted with a horizontal red dashed line.

As we previously observed for the effect of PKR variants against vaccinia K3 ^12^ deleterious PKR mutations were primarily located in the glycine-rich loop and activation loop, which are required for kinase function. Mutations to proline or stop codons were also generally deleterious (Fig. 2C). Many PKR variants had beneficial effects in the presence of variola K3 (Fig. 2C,D), indicating that the fitness landscape is not tightly constrained, in keeping with PKR’s fitness landscape against vaccinia K3. We found a strong correlation (Spearman correlation = 0.76) of PKR functional scores between the vaccinia K3 and variola K3 inhibitors (Fig. 2E). On the whole, PKR variants that evade vaccinia K3 also evade variola K3 (Extended Data Fig. 6). Similar to vaccinia K3, we found a majority of variola-K3-resistant variants are located in alpha helices D and G. We found a fitness landscape of ‘rolling hills’ in which PKR can easily access variants that evade variola K3 (Fig. 2F) rather than ‘sharp cliffs’ that would have suggested genetic variation in PKR is highly constrained. Overall, the number of PKR variants adaptive against both K3 orthologs is not much smaller than against just one or the other ortholog, despite their divergence.

### Tanapox K3

Tanapox virus is a member of the Yatapoxvirus genus, which diverged from the Orthopoxvirus genus containing vaccinia and variola virus approximately 160,000 years ago^15^. Tanapox is endemic to equatorial Africa and was first isolated from humans during an outbreak in 1957 in the Tana River Valley of Kenya^3^. It is a zoonotic virus, and while both host reservoir and mode of transmission are unknown it is speculated to have a monkey host and may be transferred to humans via infected arthropods ^4^ Tanapox K3, also known as 012 (GenBank AAD46178.1), is the most potent pseudosubstrate inhibitor of human PKR that we tested (Fig. 1B). Although Tanapox K3 shares a length of 88 residues with vaccinia K3 and variola K3, it diverges drastically at the sequence level with 39% identity to vaccinia K3 (30 identical residues) and contains both an N-terminal extension and a deletion of the basic C-terminal region predicted to interface with the PKR kinase insert (Fig. 3A). The divergence is highest in the helix insert region that contacts the PKR activation loop (Fig. 3B).

**Fig. 3.**
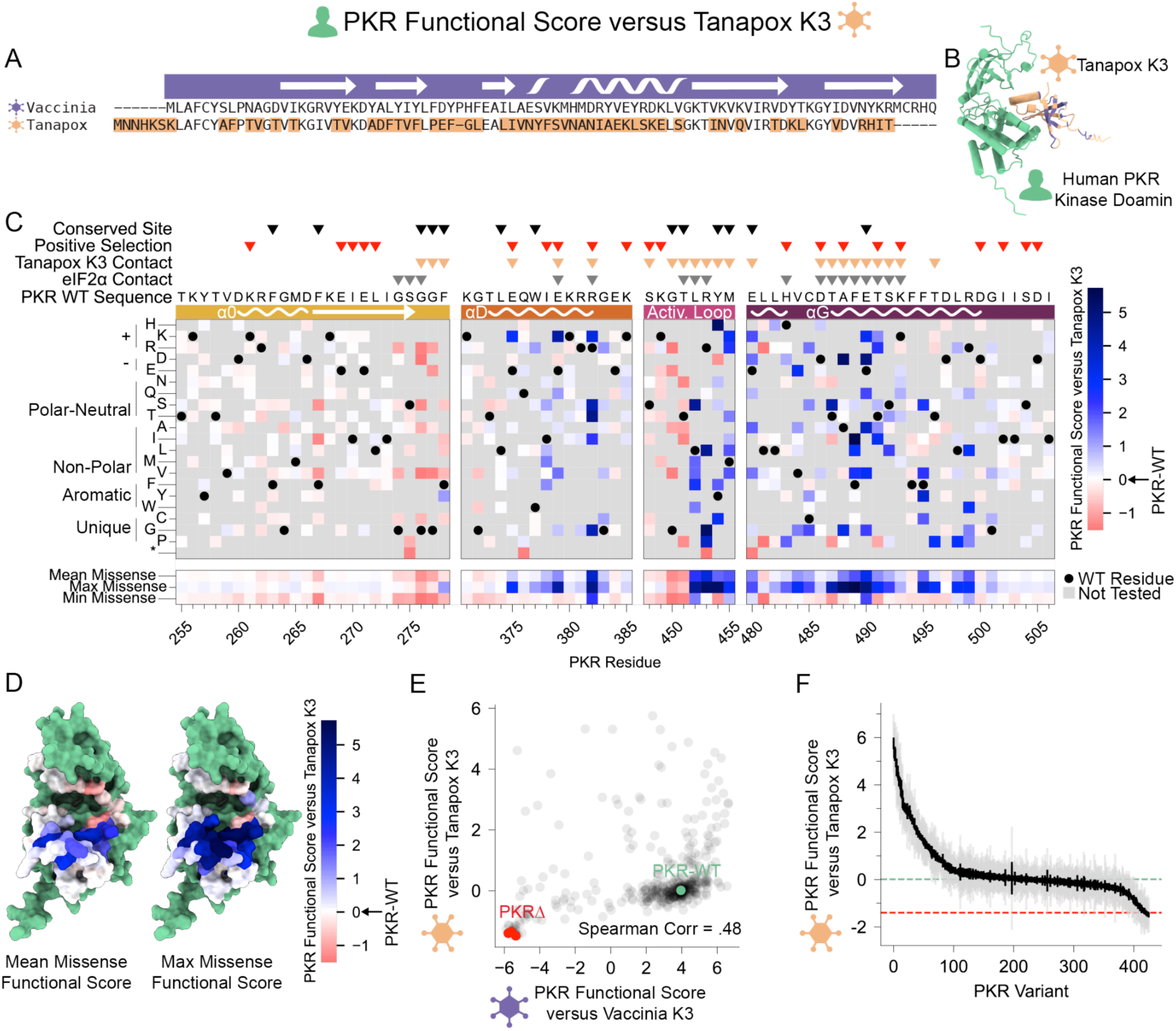
Many beneficial PKR variants are available against tanapox K3. (A) Alignment of the vaccinia and tanapox K3 homologs. (B) Prediction of tanapox K3 bound to human PKR (green), generated by AlphaFold2-Multimer. Variant sites between tanapox and vaccinia K3 are colored gold, while conserved sites are colored purple. (C) PKR functional scores versus tanapox K3 are colored ranging from susceptible (red) to WT-like (white) to resistant (blue). (D) Surface residues of the PKR kinase domain colored by their mean (left) or maximum (right) PKR functional scores against tanapox K3. (E) Correlation between PKR variant functional scores versus vaccinia and tanapox K3. (F) Ordered PKR functional scores versus tanapox K3.

Challenging our pool of PKR variants with tanapox K3 revealed that mutations outperforming wild-type PKR were widespread (Fig. 3C, D). Many of these mutations were found in helices D and G, as seen against vaccinia and variola K3, but in addition, many beneficial mutations were found in the activation loop. Given tanapox K3’s superior inhibition of PKR, we see that WT PKR and null PKR have more similar fitness in the presence of tanapox K3 than in the presence of vaccinia K3, which compresses the correlation plot of the effects of mutations against tanapox and vaccinia K3 (Fig. 3E,F). Nonetheless, we observed that the stronger inhibition of tanapox K3 did not translate to a greater ability to suppress escape, as beneficial PKR mutations were both frequent and of large effect. Moreover, beneficial mutations against vaccinia K3 again largely maintained their benefit against tanapox K3 (Extended Data Fig. 6), despite the divergence between the inhibitors (Fig. 3E). Thus, it would appear that PKR’s combined evolutionary landscape against all three of vaccinia, variola, and tanapox K3 homologs is not dramatically less navigable than against the K3 homolog from any one virus.

### *Rana catesbeiana* virus Z vIF2α

We considered two theories for the correlation between the effects of PKR mutations across the three poxvirus inhibitors. First, there may be some conserved property among the poxvirus pseudosubstrate inhibitors, shared since their common ancestor in an ancestral poxvirus, that is being exploited by the PKR mutations that confer broad resistance. Alternatively, the common property of being a PKR pseudosubstrate inhibitor may require a common feature that is exploited by the broadly resistant PKR variants. To distinguish between these two possibilities, we tested the effect of PKR mutations against an independently derived pseudosubstrate inhibitor, vIF2α from the ranavirus RCV-Z.

The Ranavirus genus, which causes systemic infections in fish, amphibians, and reptiles and has been associated with massive die-offs in these populations ^21^, is in a separate taxonomic class (*Megaviricetes*^22^) among nucleocytoplasmic large DNA viruses from the Poxviridae family. PKR is conserved in fish and amphibians, and some fish species additionally encode PKZ, an eIF2α kinase derived from PKR that is activated via Z-DNA binding domains rather than double-stranded RNA binding domains^23^ Many ranaviruses encode vIF2α, a pseudosubstrate inhibitor capable of inhibiting human PKR^6^. Ranavirus vIF2α has homology to eukaryotic eIF2α including the nucleic acid-binding OB-fold domain that interfaces with the PKR kinase domain. In contrast to the poxvirus K3 orthologs, ranavirus vIF2α is a much longer protein, and given it is an independently derived eIF2α mimic, it has very low sequence homology to vaccinia K3 (Fig. 4A).

**Fig. 4.**
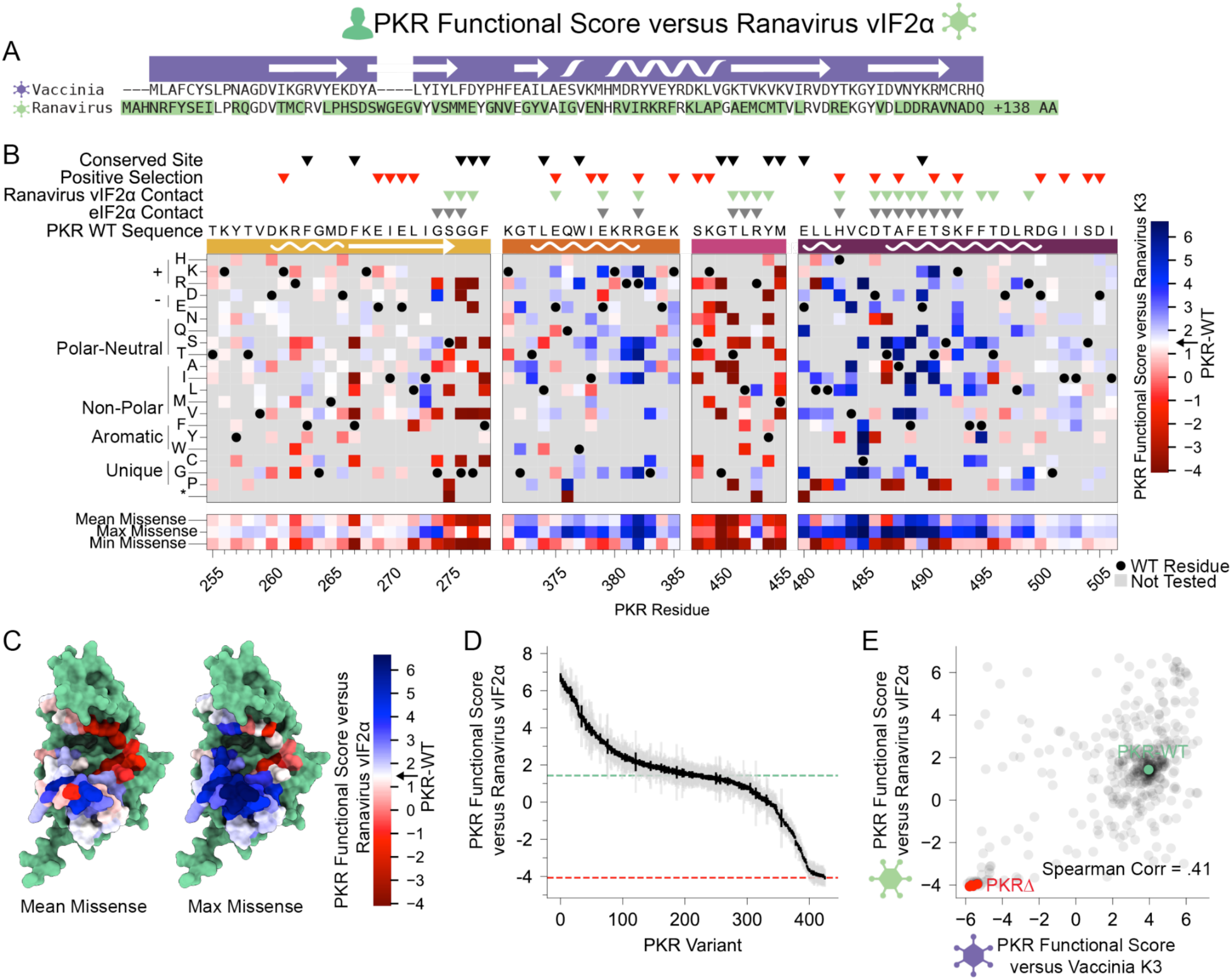
Evading PKR variants available against the independently derived Ranavirus vIF2α. (A) Alignment of ranavirus vIF2α to vaccinia K3. (B) PKR functional scores versus ranavirus vIF2α are colored ranging from susceptible (red) to WT-like (white) to resistant (blue). (C) Surface residues of PKR colored by their mean (left) or maximum (right) PKR functional scores against ranavirus vIF2α. (D) Ordered PKR functional scores versus Ranavirus vIF2α. (E) Correlation between PKR functional scores versus vaccinia K3 and Ranavirus vIF2α.

PKR mutations had a wide range of effects against ranavirus vIF2α (Fig. 4B,C). PKR functional scores versus ranavirus vIF2α exhibit a broad range with a symmetrical fitness landscape in which evading variants are abundant (Fig. 4D). Here, functional scores are less correlated with those against vaccinia K3 (Fig. 4E). However, we again see sharing of beneficial mutations, as PKR variants that increased evasion of vaccinia K3 were more likely to increase than decrease evasion of ranavirus vIF2α (46 and 15 variants, respectively) (Extended Data Fig. 6). Thus, there may be some common features of pseudosubstrate PKR inhibitors that create shared susceptibilities to a set of PKR variants, such as the need to bind PKR more strongly than eIF2α.

### Broadly evasive variants

We further examined the PKR variants that broadly evaded vaccinia K3, variola K3, tanapox K3, and ranavirus vIF2α. To normalize and combine scores across conditions we computed the z-scores for each PKR functional score distribution across each K3 condition, then computed the mean z-score across the inhibitors for each PKR variant. Plotting the mean z-score values on a heatmap again highlights α-helices D and G as harboring PKR variants that evade the viral pseudosubstrate antagonists (Fig. 5A).

**Fig. 5.**
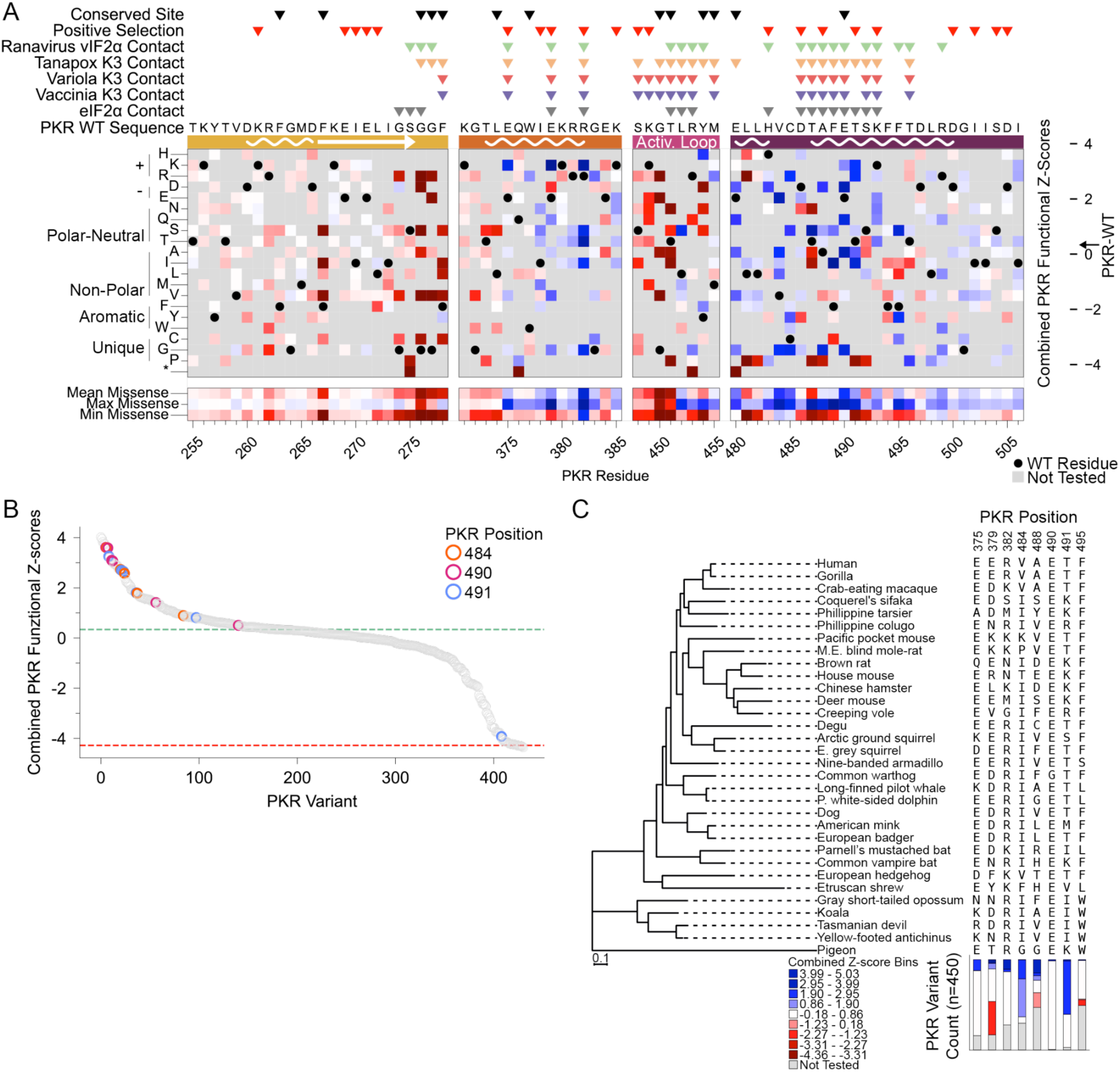
Broadly evading variants identified in circulating PKR homologs. (A) Combined PKR Functional Z-scores for vaccinia K3, variola K3, tanapox K3, and ranavirus vIF2α conditions are colored ranging from susceptible (red) to WT-like (white) to resistant (blue). (B) Ordered Combined PKR functional Z-scores highlighting variants at PKR positions 484, 490, and 491, with green and red dashed lines representing mean scores for WT PKR and nonsense variants, respectively. (C) Phylogenetic tree of select PKR homologs. Branch length corresponds to substitutions per site. Amino acid identities are shown to the right for sites at which variants of human PKR had high combined PKR functional scores. The stacked barchart below expands the analysis to 450 sequenced vertebrate species, showing the distribution of combined PKR functional scores at those sites, ranging from susceptible (red) to WT-like (white) to resistant (blue).

Interestingly, many broadly evasive PKR variants cluster to specific sites in PKR, notably PKR residues Val484, Glu490, and Thr491 (Fig. 5B). Some sites harboring broadly evasive variants were previously found to be under positive selection across vertebrates^7^, such as Thr491 and Glu379, and many of the specific variants we identify in our assay also appear in other mammalian PKR homologs (Fig. 5C). For example, isoleucine, lysine, and arginine variants of Thr491 improved evasion of all four viral inhibitors, and can be found in PKR homologs from primates, rodents, bats, and marsupials. Similarly, substituting Arg382 for lysine improved resistance to the four inhibitors, and lysine has been gained in mammals several times independently at this position. It should be noted that in other vertebrate PKR orthologs these variants exist alongside other variants, and in that context their effects may differ from the effects seen for single-residue variants of human PKR that we examined here.

Interestingly, alanine, aspartic acid, glutamine, and glycine mutations at PKR Glu490 were all broadly evasive, yet the glutamic acid residue is well conserved across vertebrate PKR homologs (Fig. 5C) and within the four human eIF2α kinases (Extended Data Fig. 7). One possibility is that the glutamic acid is required for optimal PKR function in the absence of viral inhibition, but we found that these variants of Glu490 functioned similarly to wildtype PKR (Extended Data Fig. 2). Thus, although variants of Glu490 have the potential to enable escape from viral inhibitors, we infer that some functional selection that is absent from the yeast assay has largely maintained the glutamic acid at the site across vertebrate PKR homologs.

### PKR variants with divergent effects

We also found PKR variants with differential evasion of the viral inhibitors, which could reveal interesting biological differences between the inhibitors. In many cases, the differential effects were due to differences in the inhibitors’ amino acids (Fig. 6A), particularly at contact sites with PKR. For instance, most variants of PKR-Glu375 were beneficial against vaccinia K3 and neutral against variola K3, tanapox K3, and ranavirus vIF2α (Fig. 6B). This was likely because these variants disrupt a charge interaction between the negatively charged Glu375 and vaccinia K3’s positively charged Lys45 (Fig. 6C), whereas no such benefit is expected against variola K3 Gln45, tanapox K3 Ser50, or ranavirus vIF2α Glu52. Conversely, loss of the positive charge of PKR-Arg453 was specifically deleterious against vaccinia K3 (Fig. 6B), which could be because PKR-Arg453 reduces binding by vaccinia K3 through a charge clash with Lys45 (Fig. 6C). Several variants at PKR Arg382 yielded stronger evasion of variola K3, tanapox K3, and ranavirus vIF2α than vaccinia K3 (Fig. 6B). Both Glu375 and Arg382 are on the substrate-facing surface of α-helix D (Fig. 6D) and show signatures of rapid evolution across vertebrates^7^, consistent with the optimal identity of these sites being highly dependent on the particular viral inhibitors encountered by a given species. Indeed, the viral inhibitors all have different residues that interact with Glu375 (Lys45 in vaccinia K3, Fig. 6A, D). Curiously, in contrast, Arg382 is opposite an arginine residue well conserved among the viral inhibitors and eIF2α (Arg69 in vaccinia K3, Fig. 6A, Extended Data Fig. 1), suggesting the variation at Arg382 might reflect allosteric interactions. Finally, PKR-Phe489, which protrudes from the tip of α-helix G towards the binding partners, had highly variable mutational effects (Fig. 6A). Most mutations of Phe489 dramatically improved PKR functionality in the presence of vIF2α, which could result from disruption of pi-pi interactions between Phe489 and ranavirus vIF2α’s Tyr46 (Fig. 6E). In contrast, Phe489 mutations, especially to leucine and valine, were deleterious against vaccinia and variola K3. This could indicate that the hydrophobic residues interact better with the K3 proteins’ Ile39 residues than they do with vIF2α Tyr46.

**Fig. 6.**
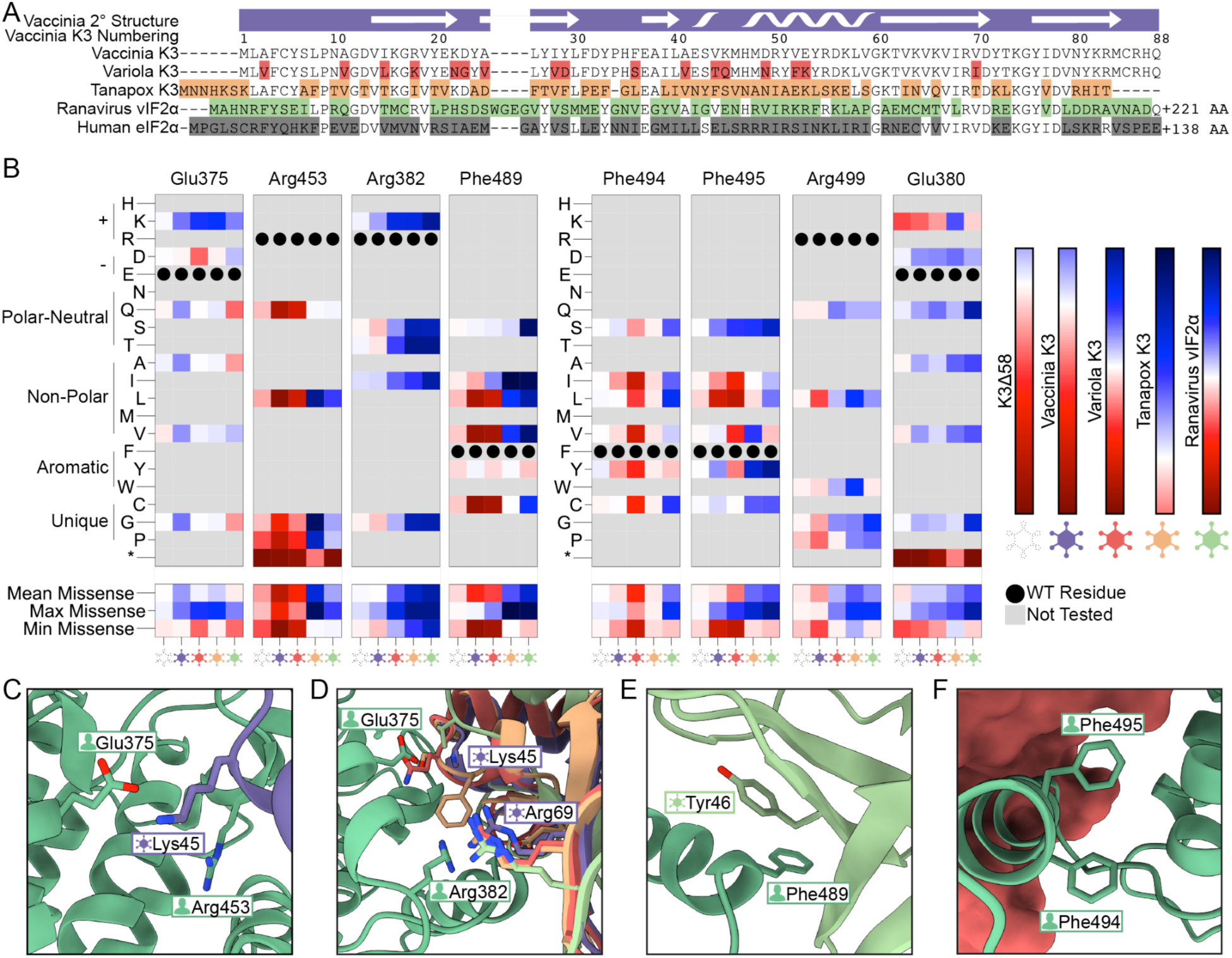
Variants at select PKR sites have divergent effects across viral pseudosubstrate inhibitors. (A) Multiple sequence alignment of viral pseudosubstrate inhibitors and human eIF2α. Residues that differ from the vaccinia K3 sequence are highlighted in red (variola K3), orange (tanapox K3), green (ranavirus vIF2α) and gray (human eIF2α). vIF2α and eIF2α sequences are not depicted after residues Gln95 and Glu94, respectively. Top tracks depict the secondary structure of vaccinia K3 and the residue numbers. Sequences were aligned using the EBI Muscle (v3.8.425) sequence alignment service. (B) PKR functional scores versus K3Δ58 (dashed line), vaccinia K3 (purple), variola K3 (red), tanapox K3 (orange), and ranavirus vIF2α (green) for select PKR sites. Cells are colored based on the prior generated maps for each pseudosubstrate condition. (C) Proximity of PKR polar residues Glu375 and Arg453 in proximity to vaccinia K3 Lys45. (D) PKR Glu375 is proximal to diverse residues across the viral pseudosubstrate proteins, as opposed to PKR Arg382 which is proximal to a fully conserved arginine residue (vaccinia K3 Arg69). (E) Pi-Pi interaction between PKR Phe489 and ranavirus vIF2α Tyr46. (F) PKR Phe494 and Phe495, located on α-helix G, form intra-molecular interactions with PKR α-helix F.

We also saw differential effects at some PKR sites whose side chains formed intramolecular interactions within PKR, rather than contacting the inhibitors. Variants at these sites may change the positioning or shape of PKR’s substrate-binding surface, which has been proposed to be a primary way for PKR mutations to evade pseudosubstrate inhibitors^8^. We observed differential effects of mutations in α-helix G of residues that face towards α-helix F (Fig. 6F). For instance, mutation of either Phe494 or Phe495 to the less bulky alanine, valine, or isoleucine were specifically deleterious in the presence of variola K3 (Fig. 6A). Variants at Arg499 were uniformly deleterious in the presence of vaccinia K3 but were beneficial against the other three inhibitors, and variants of Arg380 in α-helix D were specifically beneficial against vIF2α.

We were surprised to see that many PKR variants in residues 452-455 of the activation loop conferred strong escape specifically of tanapox K3 (Fig. 3C). The activation loop is essential for PKR’s kinase activity and has several residues conserved across PKR homologs and eIF2α kinases, including Thr451 and Tyr454 (Extended Data Fig. 7). Consistent with being functionally constrained, many activation loop variants reduced PKR function in the absence of any viral inhibitor (Extended Data Fig. 2A).

Perhaps variants with partially reduced eIF2α binding can nonetheless outperform WT PKR in the presence of a very strong inhibitor like tanapox K3, as long as they have a larger decrease in binding to the inhibitor. Consistent with this possibility, when we previously screened a hyperactive variant of vaccinia K3, K3-H47R, we noted an increase in evasive variants at the activation loop, though less pronounced than for tanapox K3^12^

### Myxoma K3

PKR appears to be readily able to acquire SNP-accessible variants that evade viral inhibitors from vaccinia, variola, tanapox, and ranavirus, including variants that evade multiple inhibitors. A separate potential limit on PKR’s evolutionary landscape is the degree to which variants render it susceptible to previously ineffective inhibitors. We screened our library of PKR variants against an additional viral inhibitor, myxoma virus M156R (hereafter referred to as myxoma K3), which does not inhibit human PKR ^24^ Myxoma virus belongs to the Leporipoxvirus genus, which diverged from the Yatapoxvirus genus that includes tanapox virus approximately 137,000 years ago and the Orthopoxvirus genus approximately 160,000 years ago^15^. Myxoma virus is native to New World rabbits, and causes the highly fatal disease myxomatosis in European rabbits.

We found the PKR functional scores versus myxoma K3 largely clustered with PKR WT (Extended Data Fig. 8A). When we compare PKR functional scores in the presence of myxoma K3 and in the absence of any PKR inhibitor (K3Δ58, Extended Data Fig. 8B) we observe a very strong correlation (Pearson correlation coefficient > 0.98). Thus, we concluded that none of the PKR variants we tested rendered PKR susceptible to myxoma K3 inhibition.

## DISCUSSION

We have expanded our characterization of human PKR’s local evolutionary space, identifying genetic variants that maintain kinase functionality when antagonized by diverse viral pseudosubstrate inhibitors that mimic PKR’s natural substrate, eIF2α. We screened single-residue variants of the kinase domain of the canonical human PKR against five viral pseudosubstrate antagonists: vaccinia K3, variola K3, tanapox K3, myxoma K3, and ranavirus vIF2α. We identified many variants that broadly evade diverse pseudosubstrate inhibitors. No variants in our library rendered PKR susceptible to myxoma K3, suggesting that PKR’s evolutionary landscape may not include substantial risk of gaining susceptibility to antagonists specialized against other species’ PKR.

Variants of PKR that evade vaccinia K3 were enriched among variants that evaded variola K3 or tanapox K3. We considered two hypotheses. First, perhaps this correlation arises from the poxviral inhibitors being related to each other, which could cause them to share susceptibility to certain variants. Second, viral pseudosubstrate mimics could share a common feature due to their common aim. Thus, we tested the effects of PKR variants against a fourth pseudosubstrate inhibitor, the independently derived vIF2α from RCV-Z. To our surprise, PKR variants that evaded vaccinia K3 were more likely to evade RCV-Z vIF2α, though the correlation was weaker than seen among the poxvirus inhibitors. To conceptualize how independently derived pseudosubstrate inhibitors would have shared susceptibilities, we formulated a model inspired by the Red Queen hypothesis. The Red Queen hypothesis posits that pairs of genes involved in evolutionary conflict constantly adapt to counter each other^10^. Perhaps in the case of PKR and a viral pseudosubstrate inhibitor, it would be easier for PKR to find useful adaptations, as it would simply need to break the interaction, whereas the inhibitor would need to find specific adaptations that restore the interaction. And if there are multiple inhibitors that use a common inhibitory mechanism, a given interaction-disrupting PKR adaptation could often translate across inhibitors, much as a modification to a lock might disrupt multiple, independently derived keys.

In support of the churn predicted by the Red Queen hypothesis, several of the broadly beneficial variant residues are present at the homologous sites in other species’ PKR proteins. We were struck, however, by the presence of broadly beneficial variants of residue 490 of human PKR, which corresponds to a glutamic acid that is generally well conserved among PKR homologs and even in other eIF2α kinases. This conservation does not appear to reflect a requirement of that glutamic acid for eIF2α kinase function, as we do not observe a decrease in the functionality of these PKR variants in the absence of viral inhibitors. We thus conclude this glutamic acid is required for some function that is not captured in our yeast assay. One possibility is that it is required for on-target specificity of PKR for eIF2α. Another possibility is that it is required for autoinhibition of PKR in the absence of dsRNA. In the yeast assay, PKR is constitutively active, possibly due to the presence of endogenous dsRNAs ^25^

We previously speculated that ideal inhibition of PKR could require the viral mimics of eIF2α to bind PKR more strongly than eIF2α does, which would necessitate some divergence from eIF2α rather than perfect mimicry. The best opportunities for such divergence could be the interfaces with functionally constrained PKR residues that are unable to explore variants that leverage the divergence from eIF2α to distinguish the mimics from the desired substrate, eIF2α. Our results may shine light on such a dynamic, in which evasive variants are available at PKR’s residue 490, but it has not been able to access them evolutionarily. We further speculated that sites on the viral inhibitors that interact with freely evolving surfaces of PKR may be less likely to diverge productively from eIF2α. For instance, eIF2α’s Arg75 is highly conserved across the viral mimics; this site interfaces with PKR’s Glu379, at which we see high diversity in vertebrates and many beneficial variants.

The viral inhibitors that we screened varied in their ability to inhibit wildtype human PKR. Tanapox virus likely has a primate host, and its inhibitor sported the strongest inhibition of PKR, suggesting that tanapox K3 might be the most adapted to inhibit primate PKRs. We considered that it could be harder for human PKR to find variants that escaped tanapox K3, if a greater binding affinity needed to be overcome. However, we saw the opposite: PKR variants that evade vaccinia K3 were generally also able to evade tanapox K3, while many variants that were neutral or even deleterious against vaccinia K3 also showed enhanced evasion of tanapox K3. We also observed that some tanapox-resistant PKR variants caused partial loss of PKR function in the absence of any inhibitor. We speculated that in the face of a very strong pseudosubstrate inhibitor, it could be beneficial to adopt variants that reduced binding to both the intended substrate and the pseudosubstrate, so long as they had a greater reduction in pseudosubstrate binding. Interestingly, a natural consequence of such adaptation could be that it would select for a later variant that ameliorated the drop in eIF2α-targeting functionality. Such compensatory evolution has previously been proposed for other viral-interacting proteins and other proteins experiencing Red Queen conflict^26,27^

We also considered the genetic variation and constraints experienced on the other side of the conflict, the viral pseudosubstrate inhibitors. Though diverse, the poxvirus pseudosubstrate inhibitors share a handful of residues that are identical to human eIF2α, notably eIF2α residues Arg75 and Lys80. The crystal structure of PKR bound to eIF2α complex^28^ (PDB 2A1A) depicts these residues forming a positively charged “pincer” that interfaces with PKR α-helices D and G, specifically Glu379, Asp486, and Glu490. As previously discussed, Glu490 is well conserved across PKR homologs, however, Glu379 and Asp486 are under positive selection ^7^. While Asp486 variants have variable effects on evading viral mimics, both Glu379 and Glu490 are broadly evading sites. The predicted structures of the viral pseudosubstrate proteins also maintain the positively charged “pincer”. This common feature appears to be a necessary constraint for the viral mimics, despite the divergence of the interfacing PKR residues. Future experiments exploring the evolutionary landscape of viral pseudosubstrate inhibitors would offer further insights into the opportunities and constraints in this genetic conflict.

## MATERIALS AND METHODS

### Identification and phylogenetic tree construction of K3 orthologs

K3 orthologs were collected using NCBI BLASTp^29^, using a vaccinia virus K3 protein sequence (NCBI YP_232916.1). K3 ortholog sequences were aligned using the EBI Muscle (v3.8.425) sequence alignment service. The output sequence alignment, phylogenetic tree, and percent identity matrix were adapted for figures in the manuscript.

### Construction of PKR and K3 ortholog plasmids

We previously constructed a base plasmid, MSp508, and inserted the human PKR allele under the inducible pGAL/CYC1 promoter to generate MSp509. The human PKR sequence was derived from p1419, a gift from Thomas Dever. A negative control plasmid was constructed with a N-terminal 465 residue deletion in the PKR-WT plasmid to form MSp614. For plasmids expressing the viral pseudosubstrate inhibitors we changed the MSp508 eukaryotic selection marker from URA3 to LEU2 and inserted the K3 ortholog under the constitutive pTDH3 promoter using a standard Gibson Assembly protocol (NEB Catalog # E2611L). The K3Δ58 and vaccinia K3 alleles were derived from MSp512 and MSp510, respectively, forming MSp515 and MSp517. Variola C3 and ranavirus vIF2α were derived from pC1800 and pC3853, gifts from Thomas Dever, to form MSp518 and MSp516, respectively. Tanapox K3^13^ and myxoma M156R^14^ plasmids, provided by Stefan Rothenburg, were used to form MSp529 and MSp530, respectively. All plasmid maps are provided as GenBank (.gb) files under the DOI: 10.5281/zenodo.19889168

### Spotting Assay

PKR and K3 plasmids were transformed into the BY4742 yeast strain using a standard lithium acetate protocol^30^, and grown on glucose selection plates lacking uracil and leucine. Duplicate colonies were picked and grown up to saturation overnight in CSM-URA-LEU (Sunrise Science Catalog # 1038-100) glucose media. Culture densities were measured at an OD600 and diluted to 1 OD/mL in YPD, followed by a 4-step 10-fold serial dilution, leaving the final 6th step blank (YPD only). Dilutions were pinned onto CSM-URA-LEU glucose and galactose plates using a replicator tool (V & P Scientific Catalog # 407AH) and incubated at 37°C for 3 days before being imaged.

### Generation of the PKR variant library

We previously generated a PKR variant library containing 426 nonsynonymous variants targeted at the kinase-substrate interface^12^. Briefly, this library was made via the design of doped primers using custom Python scripts and acquired through Integrated DNA Technologies. The primers encoded all SNP-assessible missense variants at specific residues throughout the PKR kinase domain: residues 255–278, 371–385, 448–455, and 480–506. These primers were used in PCR reactions to produce mutated fragments of the PKR gene that were then Gibson assembled into a plasmid fragment carrying the remainder of the PKR gene. Through this cloning, we also incorporated genetic barcodes downstream of the PKR gene using a pattern derived from Liu et al. ^31^ NNNNNaaNNNNNaaNNNNNttNNNNN

### PacBio sequencing of barcoded variant libraries

We identified unique genetic barcodes paired with PKR variants using the PacBio sequencing platform. Sixteen micrograms of the PKR variant library were digested with restriction enzymes AgeI-HF and NotI-HF to create a linear 2100 bp fragment. Digested fragments were size-selected using a SageELF instrument (Sage Science) before PacBio HiFi circular consensus sequencing (CCS) on a Sequel II instrument (Pacific Biosciences). From the CCS reads we generated a table of barcodes paired with PKR genetic variants using alignparse v0.2.6^32^ and custom Python scripts.

### Yeast growth assay to screen PKR variants against pseudosubstrate inhibitors

All yeast growth was at 30°C in CSM-URA-LEU, and liquid cultures were shaken at 200 revolutions per minute. To combine the PKR variant library with poxvirus K3 orthologs we performed sequential yeast transformations. We transformed each of the six K3 orthologs (K3Δ58, VACV K3, VARV C3, MYXV M156R, TPV K3, and RCV-Z vIF2α) into the yeast strain BY4742 (*MAT*α *ura3*Δ0 *leu2*Δ0 *his3*Δ *lys2*Δ0; strain name MSY2) using a standard high efficiency lithium acetate transformation protocol and plated onto 10-cm plates with 2% dextrose and Ura added back to the media (K3 plasmids have LEU selection marker).

Single colonies were picked from each plate and grown up to approximately 2 × 10^8^ cells, into which we transformed the PKR library. We plated each transformation onto a single 10-cm plate with CSM-URA-LEU glucose media, and then additionally plated a 1:1,000 dilution to estimate colony counts, which produced a minimum of approximately 60,000 colonies. Plates were split into two to form duplicate pools, and colonies were collected from each half plate by washing with 10 mL CSM-URA-LEU glucose media, resulting in 12 separate washed cultures.

To start the yeast growth assay, twelve cultures were seeded at low density (OD600 measurement of 0.01) in 40 mL media with 2% glucose and grown overnight for 16 hours. The following morning cultures were moved from 30°C to 4°C to pause growth, then restarted at an OD600 of 0.25 in 40 mL media with 2% glucose 3 hours prior to galactose induction. After 3 hours, once all cultures were in a log growth phase, approximately 1 × 10^8^ cells were taken from each culture as the starting timepoint sample (designated as “0 hours”). The cells were spun down in a 1.5 mL Eppendorf tube at 5,000 rpm for 1 minute, after which supernatant was removed and the pellet was frozen at −80°C; all subsequent timepoint samples were collected with the same method. Approximately 2 × 10^8^ cells from the remaining cultures were pelleted in 50 mL conical tubes at 3,000 rpm, supernatant was removed, and cells were resuspended in 80 mL CSM-URA media with 2% galactose (OD600 measurement of 0.125) to induce PKR expression. Cultures were incubated overnight at 30°C, 200 rpm. Additional samples were harvested at 12, 16, and 20 hours post-PKR induction, with approximately 1 × 10^8^ cells taken at each timepoint. The culture density was monitored and back diluted to maintain a log growth phase (OD600 measurement less than 1). With six K3 allele conditions (K3Δ58, vaccinia K3, variola K3, tanapox K3, myxoma K3, and ranavirus vIF2α), four timepoints (0, 12, 16, and 20 hours) and two replicates, a total of 48 samples were taken across the yeast growth assay.

### Plasmid extraction and barcode amplification

To quantify changes in barcode abundance between timepoints in the yeast growth assay, we harvested plasmids from the four timepoints for each K3 ortholog, then amplified and sequenced the PKR barcodes. Plasmids were harvested from the yeast samples using a modified QIAprep spin miniprep protocol. Sampled cell pellets of approximately 1 × 10^8^ cells were first thawed at room temperature, followed by the addition of 250 µL QIAprep P1 buffer and 2 µL zymolyase (2.5 units per µL), and incubation at 37°C for 30 minutes, followed by adding 250 µL P2 buffer and following the manufacturer instructions for the remainder of the protocol, except for eluting the plasmids using 25 uL water (Qiagen QIAprep spin miniprep, Catalog # 27106).

### PKR functional scores and screening analysis

Extracted PKR barcode sequences were amplified and sequenced on an Illumina NextSeq 1000 instrument. Barcodes extracted from the reads were mapped back to PKR variants using the table generated from the PacBio CCS HiFi reads. To generate PKR functional scores for each variant, barcodes were quantified across sample timepoints from the yeast growth assay, from which we calculated logarithmic fold-change values for each barcode relative to the 0-hour timepoint and calculated the area under the curve of fold-change values using custom Python scripts. The final PKR functional score for a given variant was taken from the mean functional score of the representative barcodes. Pearson correlation coefficients between replicate experiments for PKR variant functional scores in each K3 conditions were: 0.95 (K3Δ58), 0.77 (vaccinia K3), 0.81 (variola K3), 0.53 (tanapox K3), 0.89 (myxoma K3), and 0.97 (RCV-Z vIF2α). Replicate reads were combined for each condition, and functional scores were recalculated as described above

### Predicted PKR complexes and substrate contacts

All molecular graphics analysis were performed using UCSF ChimeraX v1.10.1 and PyMol v2.5.4. We used the AlphaFold2-Multimer (ColabFold v1.5.3) to generate structure predictions of the PKR kinase domain in complex with human eIF2α and the viral pseudosubstrates (Extended Data Fig. 9). We previously aligned the existing crystal structures of eIF2α and vaccinia K3 (PDB 2A1A and 1LUZ) to the AlphaFold2-Multimer predictions and found the predictions largely represent the crystal structures (RMSD <1). We then selected residues within 5 angstroms of the PKR kinase domain using PyMol: sele (contacts), br. (/{pdb_file_name}//A) within 5 of (/{pdb_file_name}//B) with chain ‘A’ representing the PKR kinase domain and chain ‘B’ representing the substrate protein. Predicted PKR kinase domain sites that contact eIF2α and the viral pseudosubstrate proteins are listed in Extended Data Table 1.

## Data Availability

Supplemental data, plasmid maps, structure predictions, figures, and code are available under the DOI: 10.5281/zenodo.19889168

## AUTHOR CONTRIBUTIONS

Designed research/conceptualization: M.J.C. and M.J.S. designed research; Performed research: M.J.C., T.G., and S.B.S. performed research;

Analyzed data: M.J.C., T.G., and M.J.S. analyzed data; Wrote paper: M.J.C. and M.J.S. wrote the paper.

All authors read and revised the paper.

## AUTHOR DETAILS

Michael J Chambers

- Center for Genomics and Data Science Research, National Human Genome Research Institute, National Institutes of Health, Bethesda, Maryland, USA
- Department of Computational and Systems Biology, University of Pittsburgh School of Medicine, Pittsburgh, PA, USA.

Contribution: Conceptualization, Investigation, Visualization, Writing - original draft

Competing interests: No competing interests declared

Tristan R Grieve

Center for Genomics and Data Science Research, National Human Genome Research Institute, National Institutes of Health, Bethesda, Maryland, USA

Contribution: Investigation, Analysis, Writing - review and editing

Competing interests: No competing interests declared

Sophia B Scobell

- Center for Genomics and Data Science Research, National Human Genome Research Institute, National Institutes of Health, Bethesda, Maryland, USA
- Microbiology, Virology, and Parasitology Program, Cell and Molecular Biology Graduate Group, University of Pennsylvania, Philadelphia, PA, USA.

Contribution: Investigation, Analysis, Writing - review and editing

Competing interests: No competing interests declared

Meru J Sadhu

Center for Genomics and Data Science Research, National Human Genome Research Institute, National Institutes of Health, Bethesda, Maryland, USA

Contribution: Conceptualization, Supervision, Analysis, Writing - original draft For correspondence

Competing interests: No competing interests declared ORCID: 0000-0002-8636-9102

## ACKNOWLEDGEMENTS

We thank Thomas Dever, Stefan Rothenburg, and members of the Sadhu lab for helpful discussions. We thank Thomas Dever and Stefan Rothenburg for strains and plasmids. Next-generation sequencing was performed by both the NIH Intramural Sequencing Center (NISC) and the Microarrays and Single-Cell Genomics Core of the National Human Genome Research Institute. This work utilized the computational resources provided by the NIH HPC Biowulf Cluster (http://hpc.nih.gov). This work was supported by the Intramural Research Program of the National Human Genome Research Institute, NIH (1ZIAHG200401). The contributions of NIH authors are considered Works of the US Government. The findings and conclusions presented in this article are those of the authors and do not necessarily reflect the views of the NIH or the US Department of Health and Human Services.

## Extended Data – Figures

**Extended Data Fig. 1.**
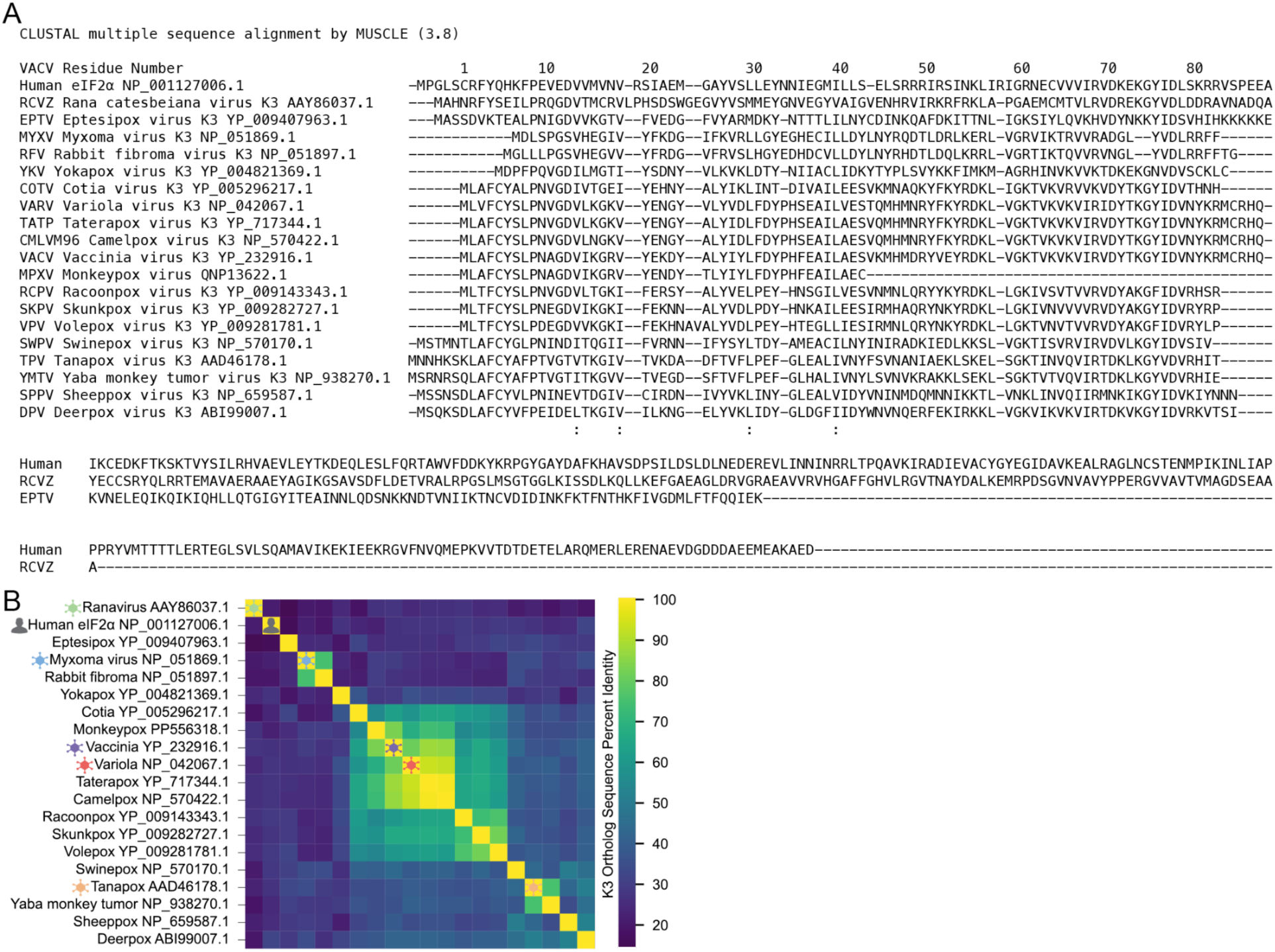
Alignment of K3 orthologs. (A) Multiple sequence alignment of viral pseudosubstrate proteins including 18 poxvirus K3 orthologs, the independently derived ranavirus vIF2α, and human eIF2α for reference. NCBI accessions are indicated for each sequence. The top track denotes vaccinia K3 residue numbers, the bottom track indicates residue conservation in Clustal format, * = fully conserved, : = strong group conservation,. = weak group conservation. Sequences were aligned using the EBI Muscle (v3.8.425). (B) A heatmap of amino acid percent identity for all-pairs of listed proteins. Five viral K3 orthologs chosen for characterization in this study are marked with virus symbols: vaccinia K3 (purple), variola K3 (red), tanapox K3 (orange), myxoma K3 (blue), and ranavirus vIF2α (green). Human eIF2α is also denoted (gray).

**Extended Data Fig. 2.**
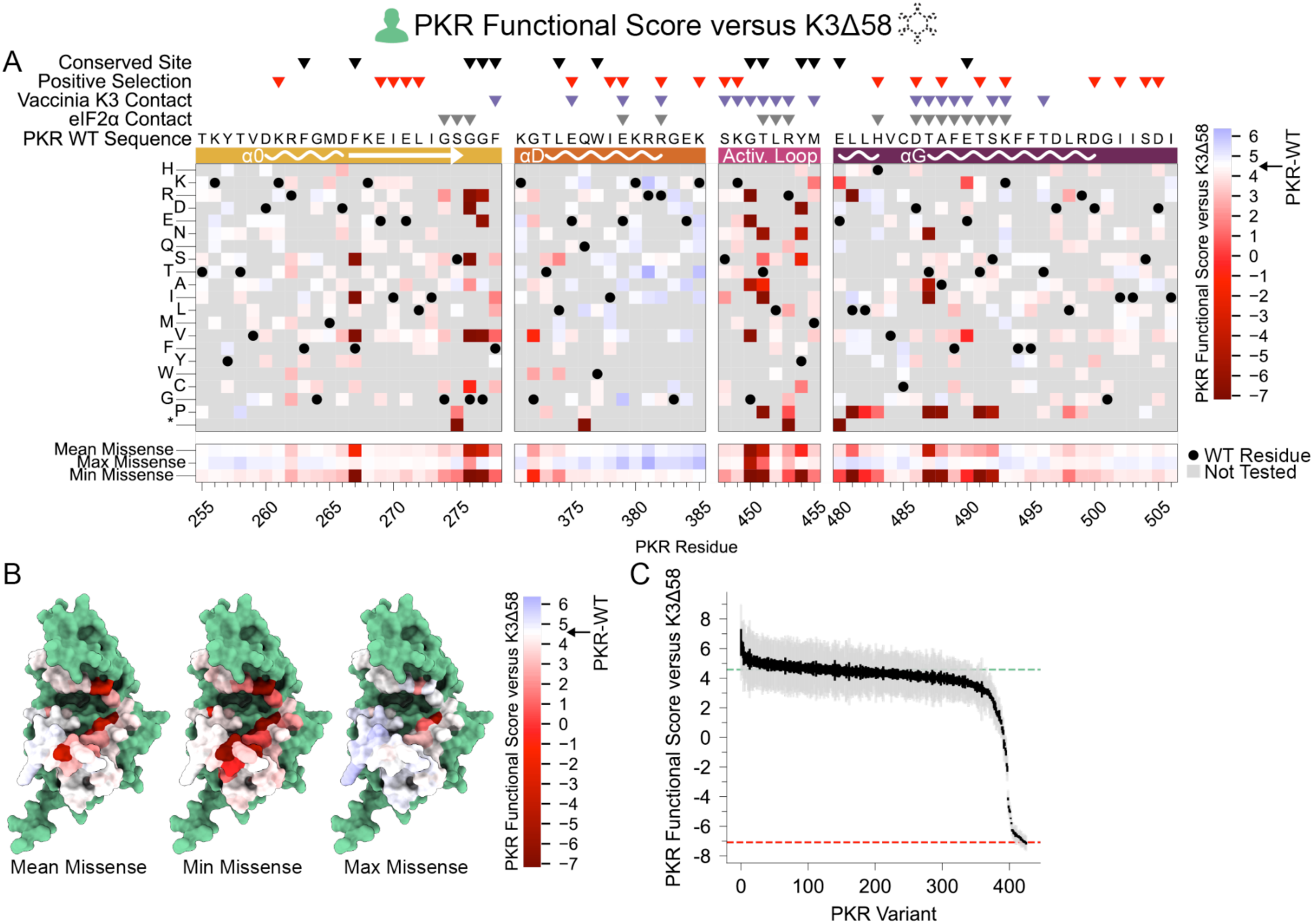
PKR functional scores versus K3Δ58. (A) PKR functional scores versus K3Δ58 are colored ranging from susceptible (red) to WT-like (white) to resistant (blue). Wild-type PKR residues are denoted with black circles. Variants not accessible through a single base change are colored in gray, as they were not characterized. Sites conserved in ≥90% of vertebrate PKR homologs or that experienced positive selection are labelled with black and red triangles, respectively. Contact sites are labelled based on AlphaFold2 prediction of being within 5 angstroms of the binding partner. (B) Surface residues of the PKR kinase domain colored by their mean (left), minimum (center), or maximum (right) PKR functional scores against K3Δ58, using identical coloring to panel A. (C) PKR functional scores versus K3Δ58, ordered by function from high to low. Gray and black lines indicate the standard deviation and standard error, respectively, in the score across the barcoded replicates of each variant. The functional score of wild-type PKR is denoted with a horizontal green dashed line, and the average functional score of the four nonsense PKR variants is denoted with a horizontal red dashed line.

**Extended Data Fig. 3.**
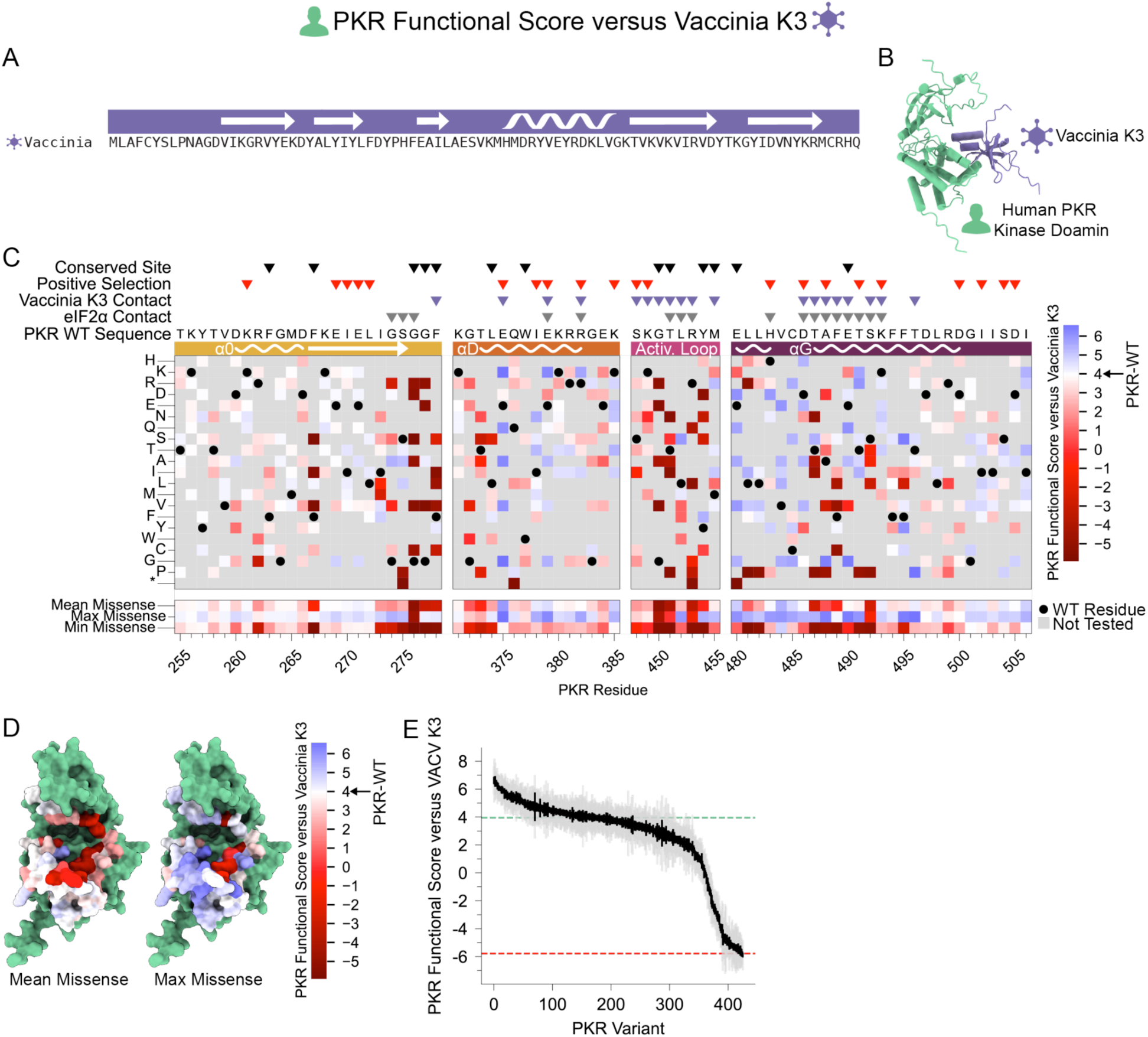
PKR functional scores versus vaccinia K3. (A) Vaccinia K3 protein sequence with secondary structure depicted above. (B) Prediction of vaccinia K3 bound to human PKR (green), generated by AlphaFold2-Multimer. (C) PKR functional scores versus vaccinia K3 are colored ranging from susceptible (red) to WT-like (white) to resistant (blue), as in Extended Data Fig. 2. (D) Surface residues of the PKR kinase domain colored by their mean (left) or maximum (right) PKR functional scores against vaccinia K3. (E) Ordered PKR functional scores versus vaccinia K3.

**Extended Data Fig. 4.**
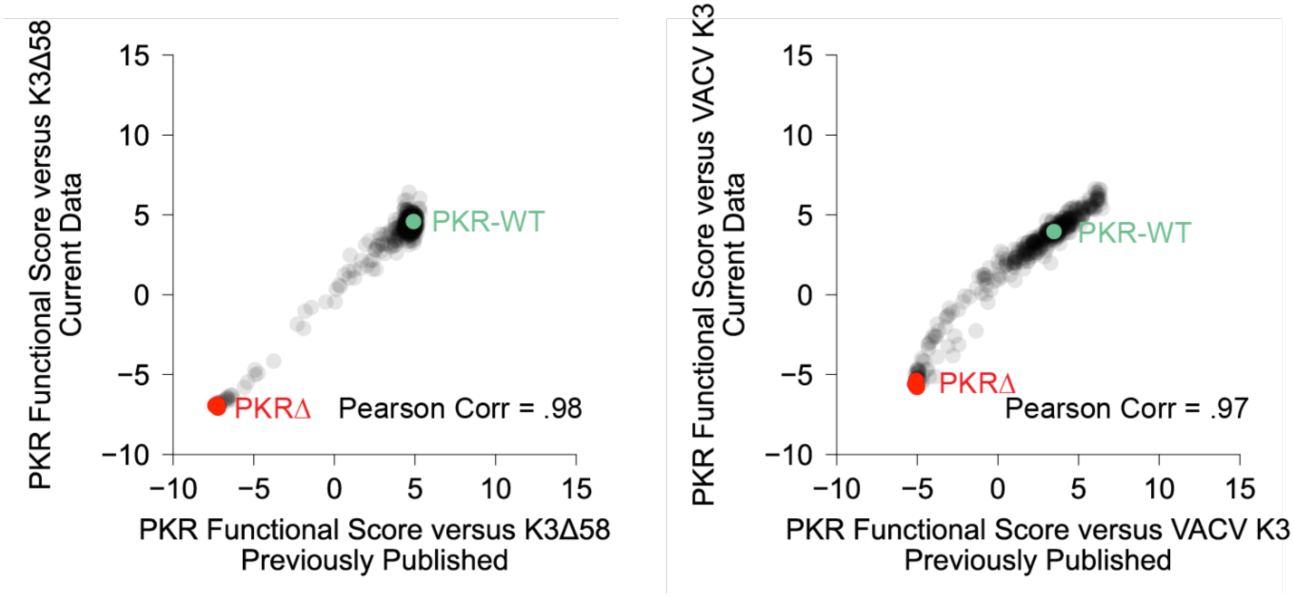
Experimental replication between current and previously published scores. PKR functional scores versus K3Δ58 (Left) and vaccinia K3 (Right). Scores for PKR WT and nonsense variants are displayed in green and red, respectively.

**Extended Data Fig. 5.**
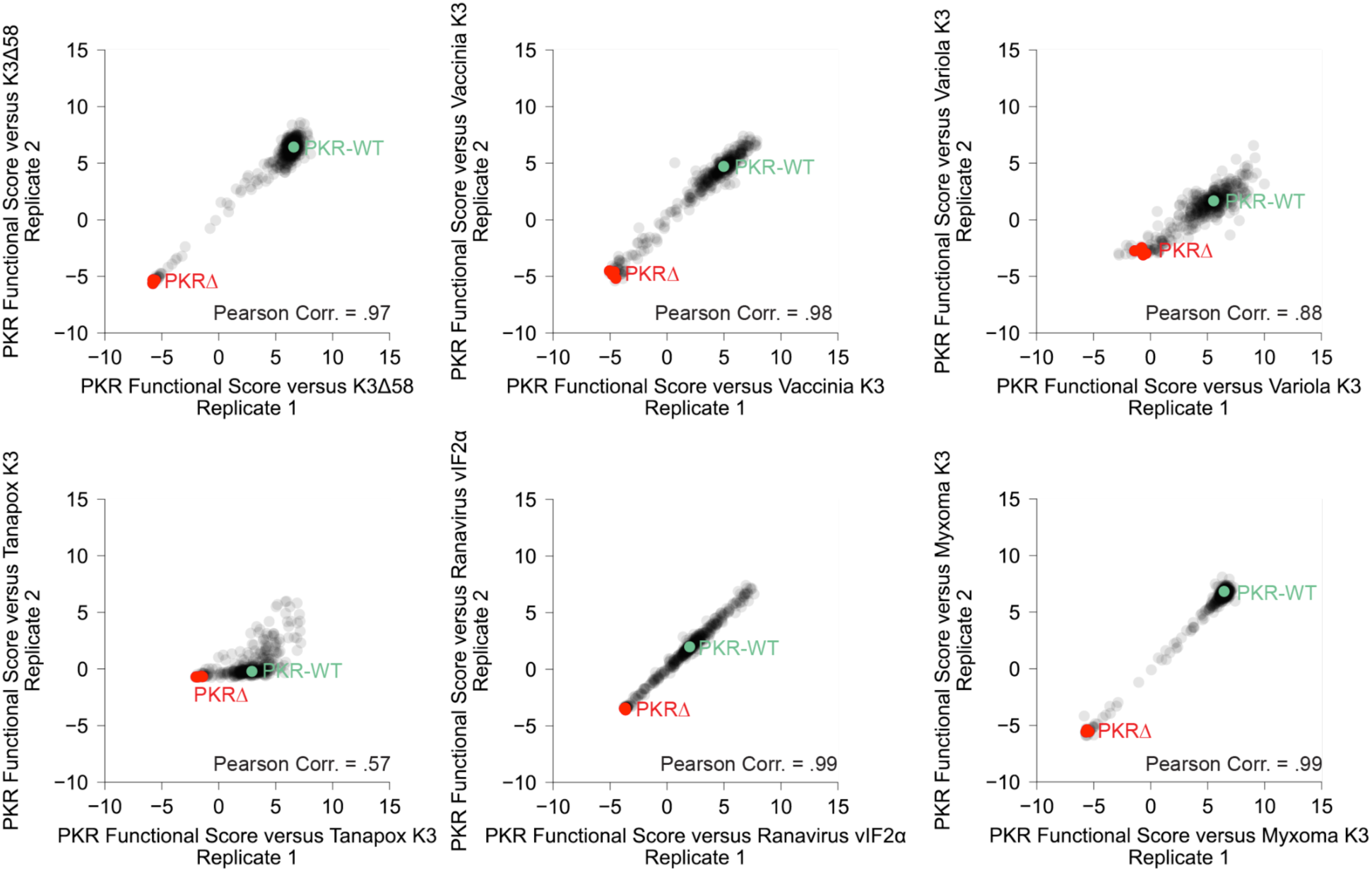
PKR functional scores for each viral pseudosubstrate condition for replicates 1 and 2. PKR WT and nonsense variant scores displayed in green and red, respectively.

**Extended Data Fig. 6.**
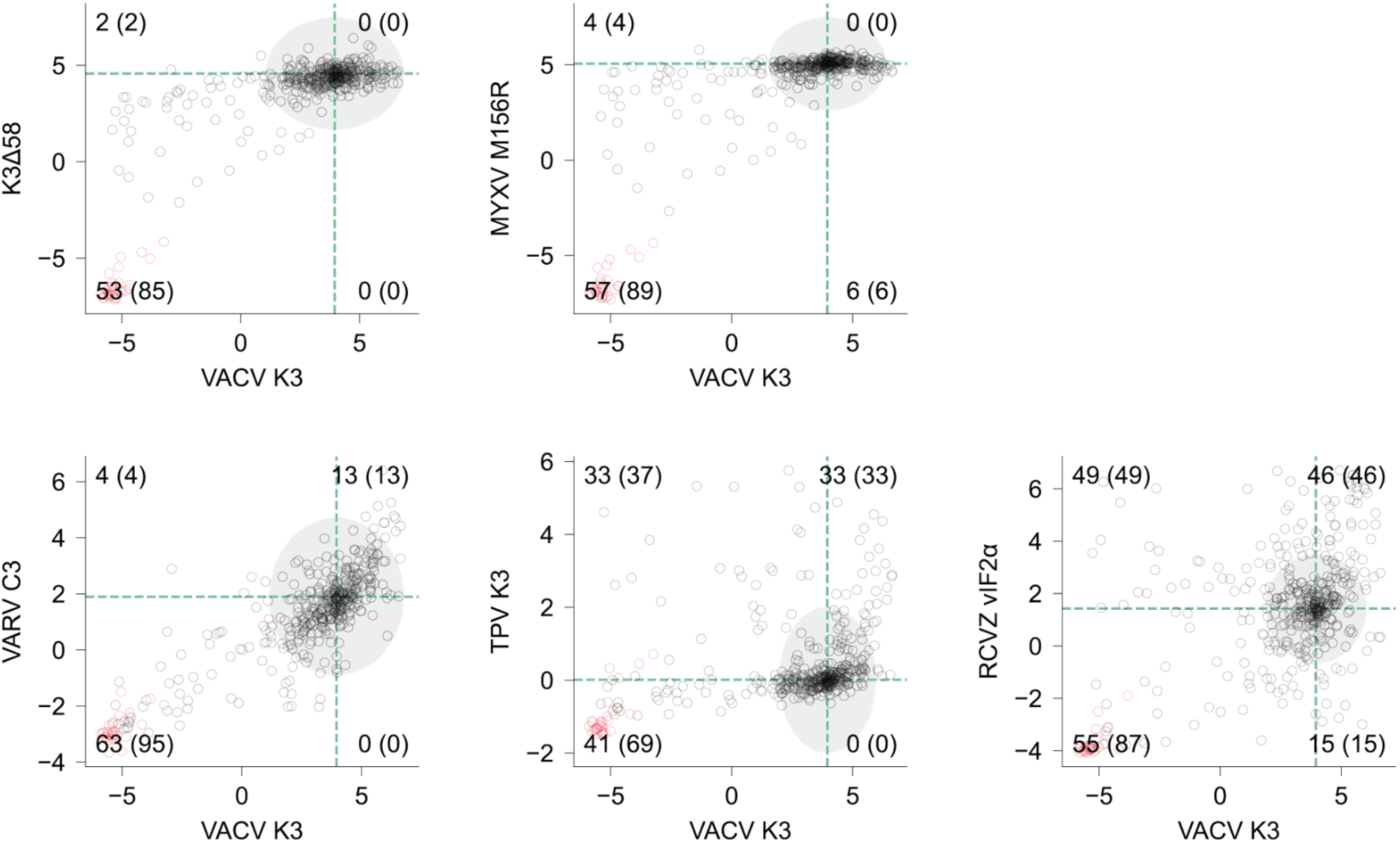
Quadrant counts for each viral inhibitor condition compared to vaccinia K3 condition. Quadrants are drawn based upon the PKR-WT functional score for each condition (green dashed lines). PKR WT and nonsense variant scores displayed in green and red, respectively. To quantify variants within each quadrant, WT-like variants falling within the PKR-WT functional score standard deviation (gray ellipse) are masked out, followed by a tally of functional variants (black circles). Quadrant tallies that include nonfunctional variants (red circles) are included in parenthesis.

**Extended Data Fig. 7.**
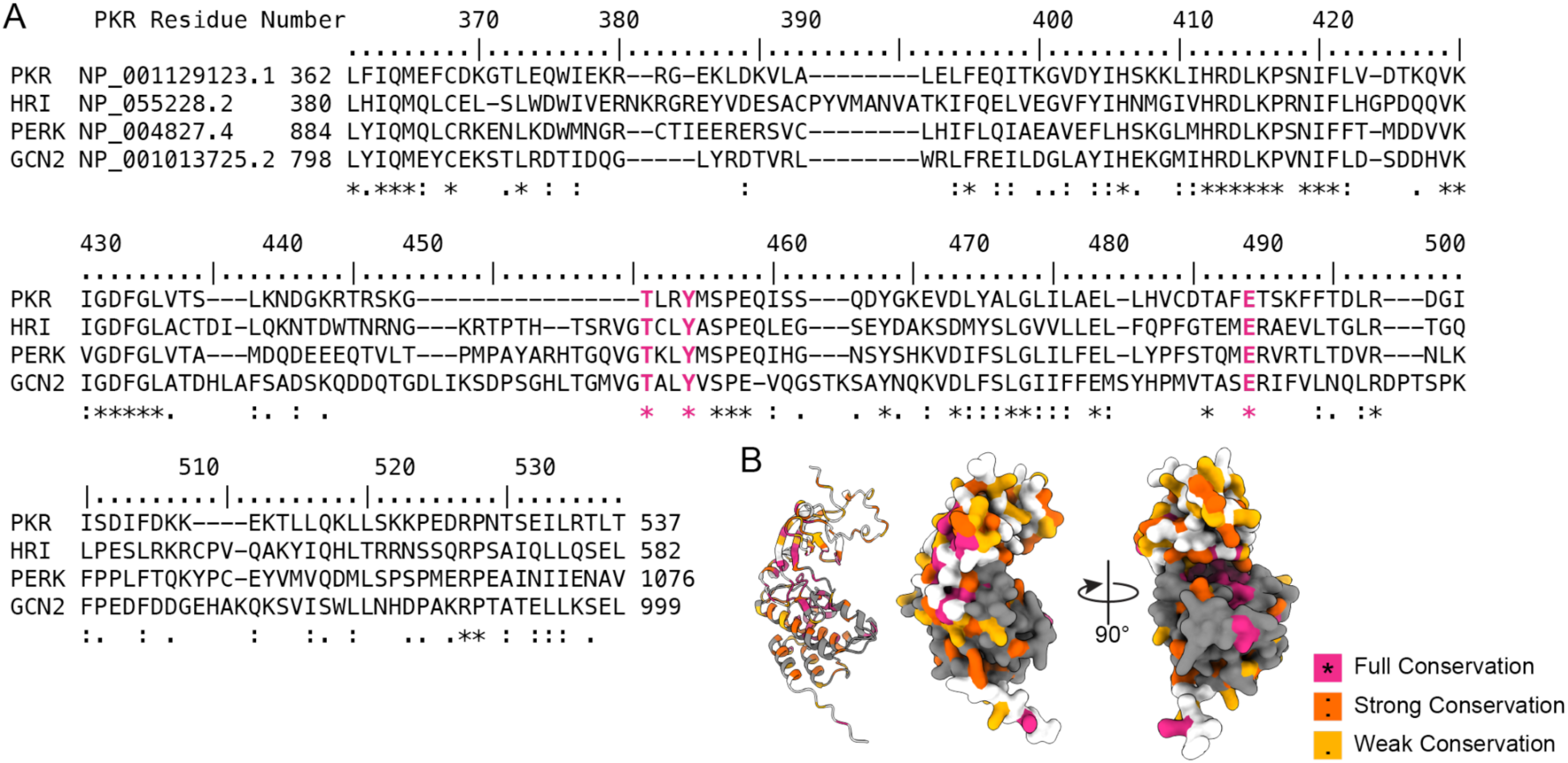
eIF2α kinase alignment. (A) Display of protein sequence alignment of all four human eIF2α kinases (PKR, HRI, PERK, and GCN2) from PKR residues 362-537. The top track denotes PKR residue number, with Thr451, Tyr454, and Glu490 highlighted in pink as referenced in the main text. Sequences were aligned using the EBI Muscle (v3.8.425). (B) AF2 structure prediction of the PKR kinase domain with residues 362-537 colored (gray), with full (pink), strong (orange), and weak (yellow) conservation residues highlighted.

**Extended Data Fig. 8.**
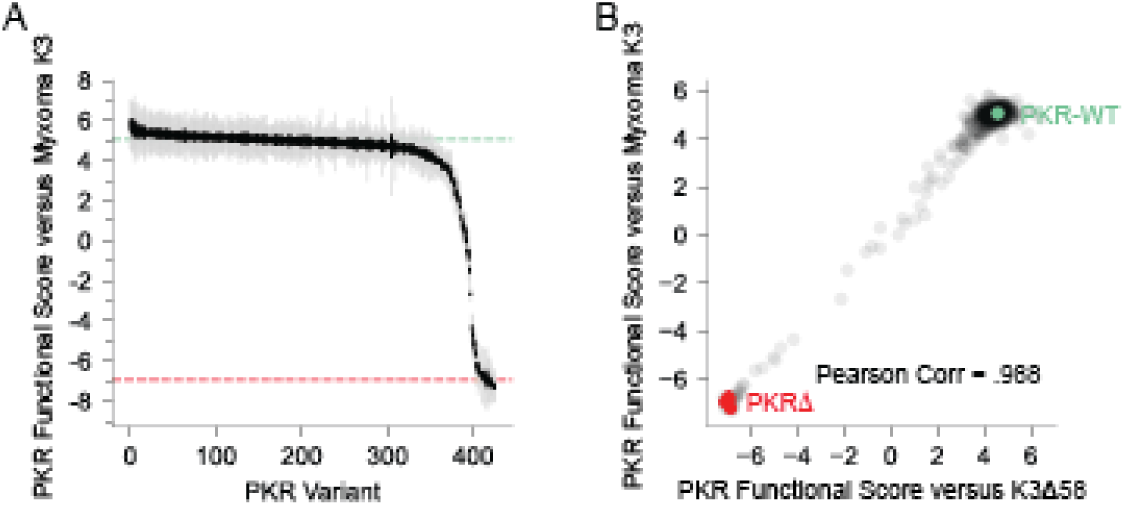
No PKR variants rendered the kinase domain susceptible to myxoma K3 inhibition. (A) Ordered PKR functional scores versus myxoma K3. (B) Correlation between PKR functional scores versus K3Δ58 and myxoma K3, with PKR-WT and nonsense variants depicted in green and red, respectively.

**Extended Data Fig. 9.**
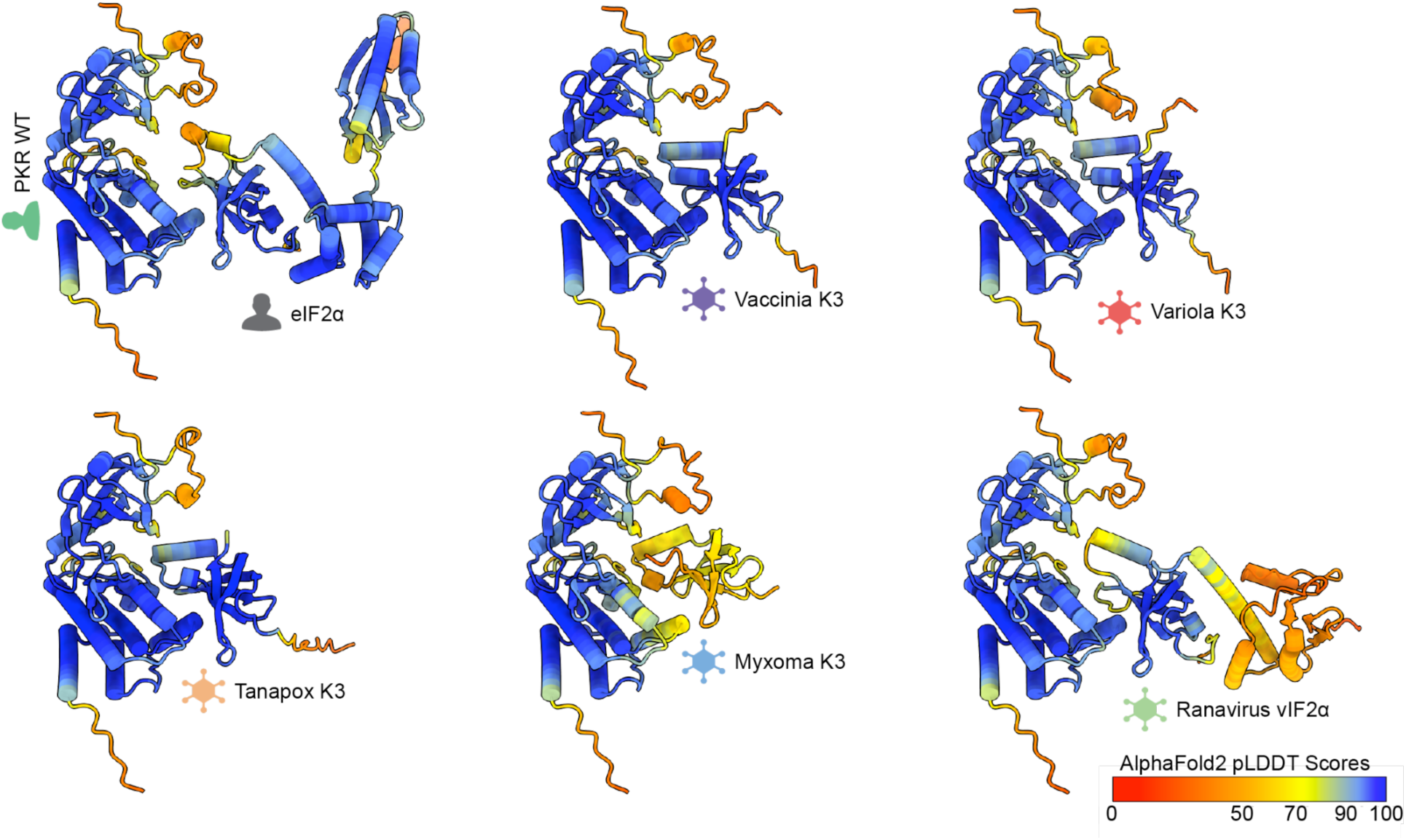
AlphaFold2-Multimer predictions for PKR kinase domains in complex with human eIF2α and viral pseudosubstrates. The PKR kinase domain (residues 250-551) were predicted in complex with each viral pseudosubstrate protein. Residues are colored by AlphaFold2 pLDDT confidence scores. Predictions were made using ColabFold v1.5.3.

## Extended Data - Tables

**Extended Data Table 1.** Predicted contact residues between the substrate (eIF2α or viral pseudosubstrates) and human PKR.

| Substrate | Predicted PKR contact residues to substrate | Predicted substrate residue contacts to human PKR |
| --- | --- | --- |
| Human eIF2α | 274,275,276,279,335,337,338,339,340,341,342,379,382,451,452,453,483,486,487,488,489,490,491,492,493 | 29,30,33,43,44,45,47,52,53,54,57,75,76,77,80,81,82,83,84 |
| Vaccinia K3 | 278,339,375,379,382,414,416,435,448,449,450,451,452,453,455,460,486,487,488,489,490,492,493,496 | 23,24,25,27,37,38,39,41,44,45,46,47,48,49,51,69,70,71,74,76,77,78,83 |
| Variola K3 | 275,276,277,278,339,340,342,343,344,345,375,379,382,414,416,417,418,435,447,448,449,450,451,452,453,454,455,460,480,486,487,488,489,490,492,493 | 23,24,25,27,37,38,39,41,43,44,45,46,47,48,49,50,51,52,54,57,69,70,71,74,76,77,78,83,85,87 |
| Tanapox K3 | 276,277,278,375,379,382,414,416,418,435,448,450,451,452,453,454,455,480,486,487,488,489,490,491,492,493,496 | 29,30,31,33,42,43,44,46,48,49,50,51,52,53,54,55,56,74,75,76,79,81,82,83 |
| Myxoma K3 | 275,277,278,304,335,337,339,375,376,414,416,434,435,448,450,451,452,453,454,455,486,487,488,489,490,492 | 19,36,37,38,39,40,41,42,43,44,47,51,55,65,67,75 |
| Ranavirus vIF2α | 275,276,277,336,337,338,339,342,375,379,382,416,451,452,453,454,483,486,487,488,489,490,492,493,495,496,499 | 27,28,29,30,31,34,44,45,46,48,51,52,53,54,55,56,76,77,78,81,82,83,84,85 |

